# Real-time analysis of single influenza virus replication complexes reveals large promoter-dependent differences in initiation dynamics

**DOI:** 10.1101/617613

**Authors:** Nicole C. Robb, Aartjan J.W. te Velthuis, Ervin Fodor, Achillefs N. Kapanidis

## Abstract

The viral RNA (vRNA) genome of influenza viruses is replicated by the RNA-dependent RNA polymerase (RNAP) via a complementary RNA (cRNA) intermediate. The vRNA promoter can adopt multiple conformations when bound by the RNAP. However, the dynamics, determinants, and biological role of these conformations are unknown; further, little is known about cRNA promoter conformations. To probe the RNA conformations adopted during initial replication, we monitored single, surface-immobilised vRNA and cRNA initiation complexes in real-time. Our results show that, while the 3’ terminus of the vRNA promoter exists in dynamic equilibrium between pre-initiation and initiation conformations, the cRNA promoter exhibited very limited dynamics. Two residues in the proximal 3’ region of the cRNA promoter (residues absent in the vRNA promoter) allowed the cRNA template strand to reach further into the active site, limiting promoter dynamics. Our results highlight promoter-dependent differences in influenza initiation mechanisms, and advance our understanding of virus replication.

## INTRODUCTION

Influenza is one of the world’s major uncontrolled diseases. Annual influenza epidemics are estimated to cause up to 650,000 deaths worldwide, while intermittent pandemics have led to the deaths of many millions of people. Influenza is caused by the influenza virus, an enveloped virus with a single-stranded, negative-sense, segmented viral RNA (vRNA) genome. Each vRNA segment is bound by the viral RNA-dependent-RNA-polymerase (RNAP) and viral nucleoprotein (NP) to form a ribonucleoprotein complex (vRNP) (1, 2). In vRNPs, the 13 nucleotides at the 5’ end and 12 nucleotides at the 3’ end of each vRNA segment are highly conserved, forming the viral promoter, which is bound by the RNAP (3, 4). The remaining part of the pseudo-circularised vRNA segment is bound by multiple copies of NP arranged in a helical, flexible rod-shape.

Replication of the vRNA is carried out by the RNAP, a multi-functional complex that is an attractive target for antiviral drug development. High-resolution structures of the RNAP have become available in recent years, providing intriguing static snapshots of how these enzymes function (3–6). The RNAP consists of three subunits: PB1 (which contains the canonical polymerase motifs typical of all template-dependent RNAPs), PA, and PB2. During an infection, vRNPs enter the host cell nucleus, where they are transcribed into mRNA by the RNAP in a primer-dependent manner. The cap-binding domain of PB2 binds 5’-capped host RNAs, which are cleaved after 10-15 nucleotides by the PA endonuclease domain, resulting in capped RNA fragments that are used as primers for transcription initiation (reviewed in 7, 8). Termination of mRNA synthesis occurs when the RNAP reaches a uridine tract located just upstream of the 5′ terminus of the vRNA, where the RNAP stutters, thereby adding a poly-A tail onto the viral mRNA (9). The same RNAP also replicates the viral genome, synthesizing a full-length, positive-sense copy of the vRNA known as complementary RNA (cRNA), which in turn serves as a template for RNAP to make more vRNA.

The structure of the RNAP-bound vRNA promoter has been resolved by x-ray crystallography (Fig. S1A); in the structure, the 5′ proximal region (nucleotides 1–10) forms an intramolecular stem-loop that is tightly bound in a pocket formed between PA and PB1 (3, 4, 6). Binding of the 5’ end results in ordering of the polymerase active site and activates multiple polymerase functions (10–12). The 5′ distal region (nucleotides 11-14/15) base-pairs with the 3’ distal region (nucleotides 10-13/14) to form a short duplex. The 3′ proximal region (nucleotides 1-9) can adopt two different conformations, either a pre-initiation state in which the 3’ vRNA is bound in an arc shape on the outside of the protein (4), or an initiation state in which the 3’ terminus of the vRNA threads through the template channel into the RNAP active site (Fig. S1B) (6, 13). The structure of the closely related La Crosse bunyavirus RNAP shows the proximal 3’ RNA in a third possible conformation, where the 3’ end is tightly and specifically bound in an extended, single-stranded configuration in a narrow cleft on the outside of the protein (10). The existence of alternative conformations where the proximal 3’ vRNA does not access the template tunnel suggests that off-pathway structural states may exist, and that the 3’ end must translocate into the active site before replication is initiated; however, no information on these dynamic states is available. Once the vRNA 3’ end enters the active site, replication initiates, without the need for a primer, at positions 1 and 2 of the 3’ vRNA terminus (14). It has been shown that the priming loop, a β-hairpin in the PB1 subunit that protrudes towards the active site of the RNAP (3, 4), acts as a stacking platform for the initiating nucleotide during terminal initiation on the 3′ vRNA (15).

Similarly to the vRNA, the 5’ and 3’ ends of each cRNA segment are conserved and form the cRNA promoter. As cRNAs are produced, they are bound by newly-made RNAP and NP to form cRNP complexes, which in turn synthesize new vRNAs (reviewed in 7, 8). Only limited structural information on the conformation of the cRNA promoter is available; however, it is known that the conformation of the proximal 5′ cRNA (nucleotides 1-12) is virtually identical to that of the proximal vRNA 5′ terminus (12), suggesting that the 3′ end of the cRNA promoter may adopt a similar conformation to the vRNA promoter (Fig. S1C). However, unlike initiation from the vRNA promoter, the priming loop is dispensable for internal initiation on the cRNA promoter (15). Instead, initiation on the cRNA promoter is suggested to use a ‘prime-and-realign’ mechanism, which starts internally at positions 4 and 5 to produce pppApG, followed by realignment of the pppApG product to bases 1 and 2 of the cRNA 3’ terminus, before elongation (14, 16). The proximal 3’ end of the cRNA promoter must therefore be positioned differently in the active site to the vRNA 3’ end, resulting in nucleotides 4 and 5 being positioned correctly for internal initiation (Fig. S1C).

In this study, we have used single-molecule Förster resonance energy transfer (smFRET) to study the conformations and dynamics of the influenza virus vRNA and cRNA promoters during replication initiation. Building upon our previous solution-based smFRET studies (13, 17), we performed real-time studies of surface-immobilised initiation complexes for the first time, and showed that the 3’ vRNA in individual replication complexes exists in dynamic equilibrium between the pre-initiation and initiation conformations, on the 1-second timescale. In contrast, the cRNA promoter showed virtually no dynamics, strongly suggesting that there are substantial differences in the initiation mechanisms for the two promoters. We also show that structural differences in the cRNA promoter, resulting in different positioning of the proximal 3’ cRNA compared to 3’ vRNA in the RNAP active site, are responsible for stabilising promoter binding by the RNAP. Our results offer fresh insight into the differences between initiation mechanisms for the vRNA and cRNA promoters. Our techniques and findings are also applicable to many other polymerase-nucleic acid interactions.

## MATERIALS AND METHODS

### RNA and protein

RNAs corresponding to the 3’ and 5’ conserved ends of the neuraminidase vRNA or cRNA gene segments were custom synthesized and labelled with Cy3B and ATTO647N by IBA (Gottingen, Germany). The single-stranded RNAs were annealed in annealing buffer (50 mM Tris-HCl pH 8, 1 mM EDTA, 500 mM NaCl). RNA sequences are as follows: 5’ vRNA 5’-AGUAGUAACAAGGAGUUXAAA-3 where X=U-Cy3B, 3’ vRNA 5’-UUUAAACUCCUGCUUUUGCX-3’ where X=U-Atto647N, 3’ vRNA 5’-UUUAAACUCCUGCUUUXGCU-3’ where X=U-Atto647N, 5’ cRNA 5’-AGCAAAAGCAGGAGUUX-3’ where X=U-Cy3B and 3’ cRNA 5’-AAACUCCUUGUUUCUACX-3’ where X=U-Atto647N. Recombinant influenza A/WSN/33 (H1N1) histidine-tagged RNAP (with a deca-histidine tag and a protein-A tag on the C-terminus of the PB2 subunit) was expressed in mammalian cells and purified using IgG-sepharose as previously described (18).

In vitro assays. An in vitro ApG primed replication assay was used to show that the decahistidine tag had no effect on protein activity, as previously described (13). RNP reconstitutions and primer extensions were carried out as previously described (19, 20).

### Sample Preparation

RNAP-promoter RNA replication complexes were formed by incubating 10 nM dsRNA with influenza RNAP in binding buffer (50 mM Tris-HCl (pH 8), 500 mM NaCl, 10 mM MgCl_2_, 100 µg/µl BSA, 1 mM DTT and 5% glycerol) for 15 min at 24 °C before the complexes were diluted to a final RNA concentration of 0.5 nM for imaging. Neutravidin-coated glass coverslip chambers were prepared as described (21). A 10 nM solution of biotinylated penta-His antibody (Qiagen) was incubated for 10 min on the neutravidin-coated surface and excess unbound antibody removed (22, 23). RNAP-promoter RNA complexes were incubated on the slide surface for 1 min followed by washing to remove excess unbound complexes. Imaging buffer containing 50 mM Tris-HCl (pH 8), 500 mM NaCl, 10 mM MgCl_2_, 100 µg/µl BSA, 1 mM DTT, 5% glycerol, 2 mM UV-treated Trolox and an enzymatic oxygen scavenging system consisting of 1 mg.mL^−1^ glucose oxidase, 40 μg.mL^−1^ catalase, and 1% (wt/vol) glucose was added to the observation chamber prior to imaging. Experiments were performed at 22°C unless indicated otherwise. Solution-based confocal experiments were carried out as previously described (13, 17).

### Instrumentation

Single-molecule total internal reflection fluorescence (TIRF) experiments were performed on a custom-built objective type TIRF microscope, as previously described (24, 25). Green (532-nm Cobolt Samba) and a red (635-nm Cube Coherent) lasers were alternated using alternating laser excitation (ALEX), and data were acquired using either a 100-Hz or 10-Hz alternation rate (indicated where appropriate). Excitation powers of 3 mW (green) and 1 mW (red) were used for 100-Hz experiments, while excitation powers of 1 mW (green) and 0.25 mW (red) were used for 10-Hz experiments. Solution-based confocal experiments were carried out as previously described (13, 17).

### Data Analysis

Fluorescence intensities were extracted from images using previously described TwoTone software (26), and the apparent FRET efficiency, E*, was calculated as described (27). Molecules were selected for analysis using well-defined criteria (24, 25): i) Only molecules detected in both emission channels were selected (with the channel filter ‘DexDem&&AexAem&&DexAem’, i.e. colocalization of the donor dye signal upon donor laser excitation, the acceptor dye signal upon acceptor laser excitation, and the acceptor dye signal upon donor laser excitation); ii) a width limit between the donor and acceptor of between 1 and 2 pixels was set; iii) a nearest-neighbour limit of 6 pixels was set and iv) a maximum ellipticity of 0.6 was used to exclude very asymmetric PSFs. We then manually inspected intensity time trajectories and selected molecules for further analysis using the following criteria: i) bleaching in a single step; ii) no donor photoblinking; iii) no acceptor photoblinking; iv) fluorescence intensities being within a limited range and v) no prominent defocusing.

Histograms of E*, or kymographs showing multiple FRET traces in one field of view, were constructed using selected molecules. We also manually classified the intensity time trajectories of individual molecules into groups. Static complexes were defined as time trajectories showing a stable E* signal with no fluctuations to higher or lower FRET states during our observation period. Dynamic complexes were defined as time-trajectories showing anti-correlated changes in the DD and DA intensities during the observation period (50 seconds).

### Hidden Markov Modelling (HMM) Analysis

HMM analysis was performed on time-trajectories displaying dynamic FRET fluctuations using a modified version of the ebFRET software (28, 29). Each time trace was fitted with 2 states, corresponding to the initiation and pre-initiation states. The dwell times for each state category were extracted and used to generate dwell-time distributions, which were fitted with an exponential decay curve to extract the mean dwell times of both states. Binarised plots, where the HMM states were colour-coded and the traces sorted by trace length, were used to give an overview of all traces in an experiment.

### Modelling of the influenza virus RNA and FRET positioning and screening software (FPS)

A model which incorporated the influenza B polymerase (PDB code 5MSG), residues 1–14 of the 5’ RNA (PDB code 5MSG), residues 1–6 of the 3’ RNA in the pre-initiation state (PDB code 4WRT) and residues 1-13 of the 3’ RNA in the initiation state (PDB code 5MSG) was created in PyMol, and the duplex RNA was extended to positions 18 for the 5΄ and 17 for the 3΄ RNA using the DuplexFold module of the RNAstructure webserver (30) and the Chimera plugin Assemble2 (31). This structure was used to model the positions of the donor and acceptor dyes. The attachment point for each dye was identified (a uridine on the C5΄ of the sugar residue). The Cy3 dye was characterized by a linker length of 14.2 Å, a linker width of 4.5 Å and dye radii of 8.2, 3.3 and 2.2 Å (x, y and z, respectively) (17). The ATTO647N dye was characterized by a linker length of 17.8 Å, a linker width of 4.5 Å and dye radii of 7.4, 4.8 and 2.6 Å (17). The accessible volumes of the dyes attached to the DNA were calculated using FRET positioning and screening software (FPS) (32, 33). Here, the sterically accessible positions of the dye were obtained from lattice calculations, with the dye modelled as a sphere. The accessible volumes were approximated by a 3D Gaussian, the centre of which was given as the average position of the dyes (represented as spheres in the figures).

## RESULTS

### The 3’ vRNA in immobilised replication complexes is highly dynamic

Crystal structures of the influenza RNAP (4, 6), as well as our previous biophysical characterization (13), revealed that the proximal 3’ end of the vRNA promoter exists in two different conformations upon RNAP binding, corresponding to pre-initiation and initiation states (Fig. 1A). In order to test whether these two states exist independently or interconvert within individual molecules, and to recover the timescales of any dynamics, we developed a total-internal reflection fluorescence (TIRF) microscopy assay for real-time observations of immobilized RNAP-promoter vRNA complexes for extended periods. The complexes were immobilized via the C-terminus of the PB2 subunit of the RNAP using a decahistidine fusion tag and biotinylated anti-histidine antibody (Fig. 1A) (24, 25). An in vitro activity assay confirmed that the decahistidine tag had no effect on RNAP activity (Fig. S2).

**Figure 1.**
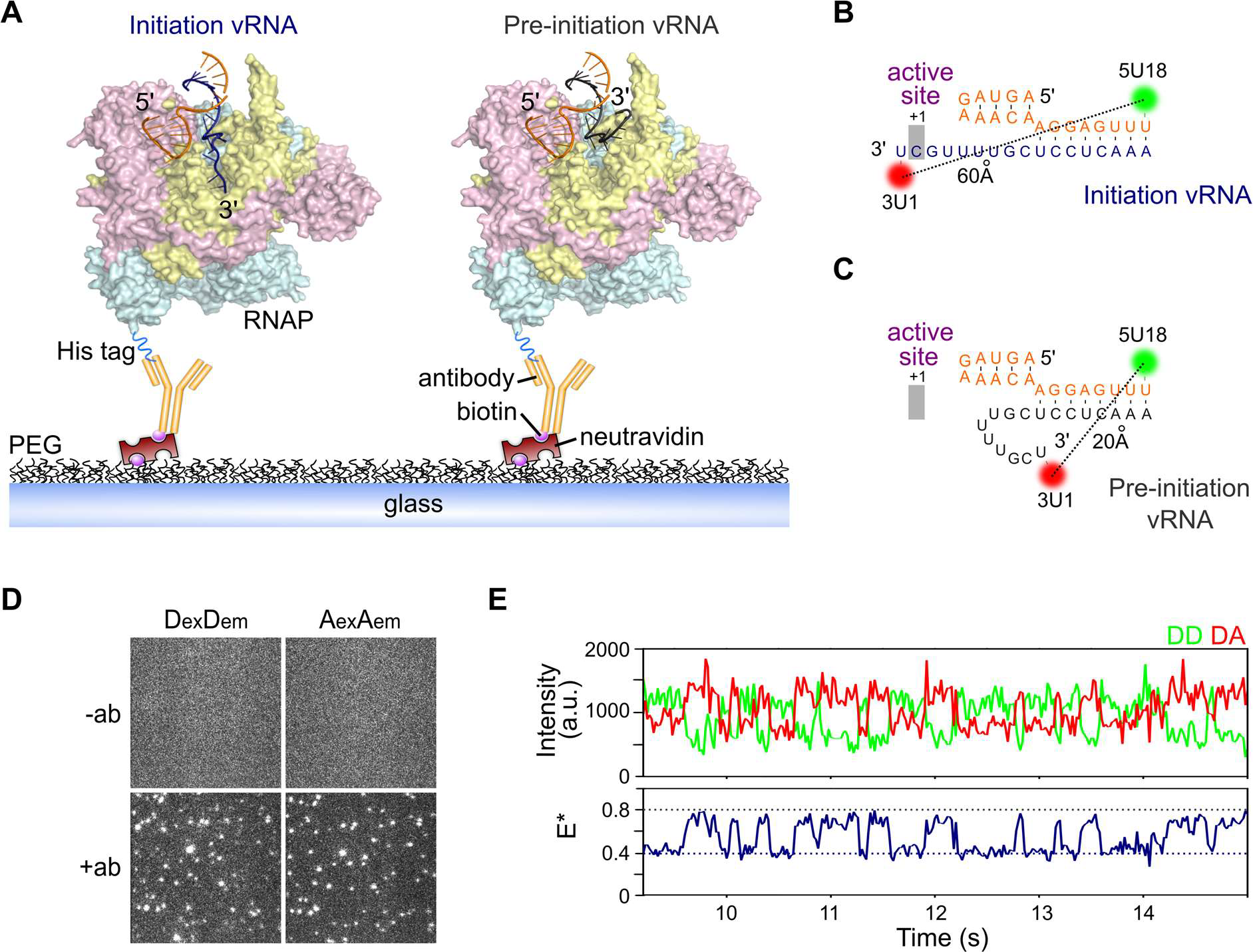
The vRNA promoter within immobilized influenza replication complexes can adopt multiple conformations. A) Cut-away structural model of the influenza RNAP bound to the vRNA in an initiation state where the 3’ vRNA (PDB code 5MSG) enters the RNAP active site (left), or bound to the vRNA in a pre-initiation state (PDB code 4WRT), where the 3’ vRNA is located on the outer surface of the RNAP (right). The RNAP (PDB code 5MSG) is shown in yellow (PB1), cyan (PB2) and pink (PA), with the 5’ vRNA in orange and 3’ vRNA in dark blue (initiation) or grey (pre-initiation). The RNAP is immobilized on a PEG-coated surface via a biotinylated anti-histidine antibody against a His_10_ tag on the C-terminus of the PB2 subunit. B&C) Fluorophore positions and sequences of the vRNA promoter used in immobilized complexes. D) Representative field-of-view of the donor (DexDem) and acceptor (AexAem) channels, showing that immobilization of complexes is specific to the anti-histidine antibody. E) Example time trace showing FRET from an immobilized replication complex containing promoter vRNA labelled with donor and acceptor fluorophores at positions 18 on the 5’ end and 1 on the 3’ end. Clear donor (DD)-acceptor (AA) anti-correlations are characteristic of FRET dynamics. Frame time: 20ms. E* represents apparent FRET efficiency.

Prior to surface immobilisation, the RNAP was incubated with short, partially complementary RNAs corresponding to the conserved 5’ and 3’ termini of the vRNA, fluorescently labelled with a donor dye at position 18 on the 5’ strand (5U18) and an acceptor dye at position 1 on the 3’ strand (3U1) (Fig. 1B&C). We have shown previously that labelled RNAs are bound by the RNAP with high affinity (17), and that these positions do not affect the activity of the RNAP in an in vitro replication assay (13). Dyes at these positions are also able to report on the conformational state of the 3’ vRNA, as we have demonstrated using precise modelling of the dye positions on structural models of the vRNA promoter bound to RNAP (Fig. S3A) and previously with a solution-based smFRET assay (13, 17).

In the initiation state, the dyes are far apart (~60Å, Fig. 1B), corresponding to a low-FRET value, while in the pre-initiation state, the dyes are in close proximity (~20Å apart), corresponding to a high-FRET value (Fig. 1C) (13). In the absence of antibody, the non-specific absorption of free promoter RNA to the surface was negligible (Fig. 1D). Following the immobilization of replication complexes, we performed TIRF measurements, and used the donor and acceptor fluorescence signals to calculate apparent FRET efficiencies (E*). An initial visual inspection of time-traces revealed that the 3’ vRNA within individual replication complexes was highly dynamic, exhibiting clear transitions between high (E*~0.8) and low (E*~0.4) FRET states (Fig. 1E).

To further characterize the conformational dynamics of the 3’ vRNA, we inspected individual time-traces of many (n=106) immobilized RNAP-vRNA promoter complexes containing fluorophores at position 18 on the 5’ strand, and position 1 on the 3’ strand (Fig. 2A and S4A). To study all replication complexes in a single field of view, we superimposed their FRET traces onto a “kymograph” (Fig. 2B). We also depicted the traces as FRET histograms, which were fitted with Gaussian functions to determine the mean FRET efficiency of the distributions (Fig. 2C). Replication complexes in the absence of nucleotides resulted in a bimodal FRET distribution centered at E*~0.44 and E*~0.73 (Fig. 2C). Based on our dye modelling and previous smFRET measurements (13), we assigned the high-FRET state to vRNA in the pre-initiation state, and the low-FRET state to vRNA in the initiation state. The large majority of complexes (87%) showed dynamics, exhibiting rapid transitions between high- and low-FRET states (e.g., Fig. 2D), while the remaining molecules showed either stable low FRET (11%) or stable high FRET (2%). Increasing the temperature from 22°C to 37°C produced a single FRET population centered at E*~0.53 (Fig. S5A-D), suggesting that vRNA dynamics likely occur on a timescale faster than our temporal resolution, a finding which was further supported by confocal data taken at 37°C (Fig. S5E&F).

**Figure 2.**
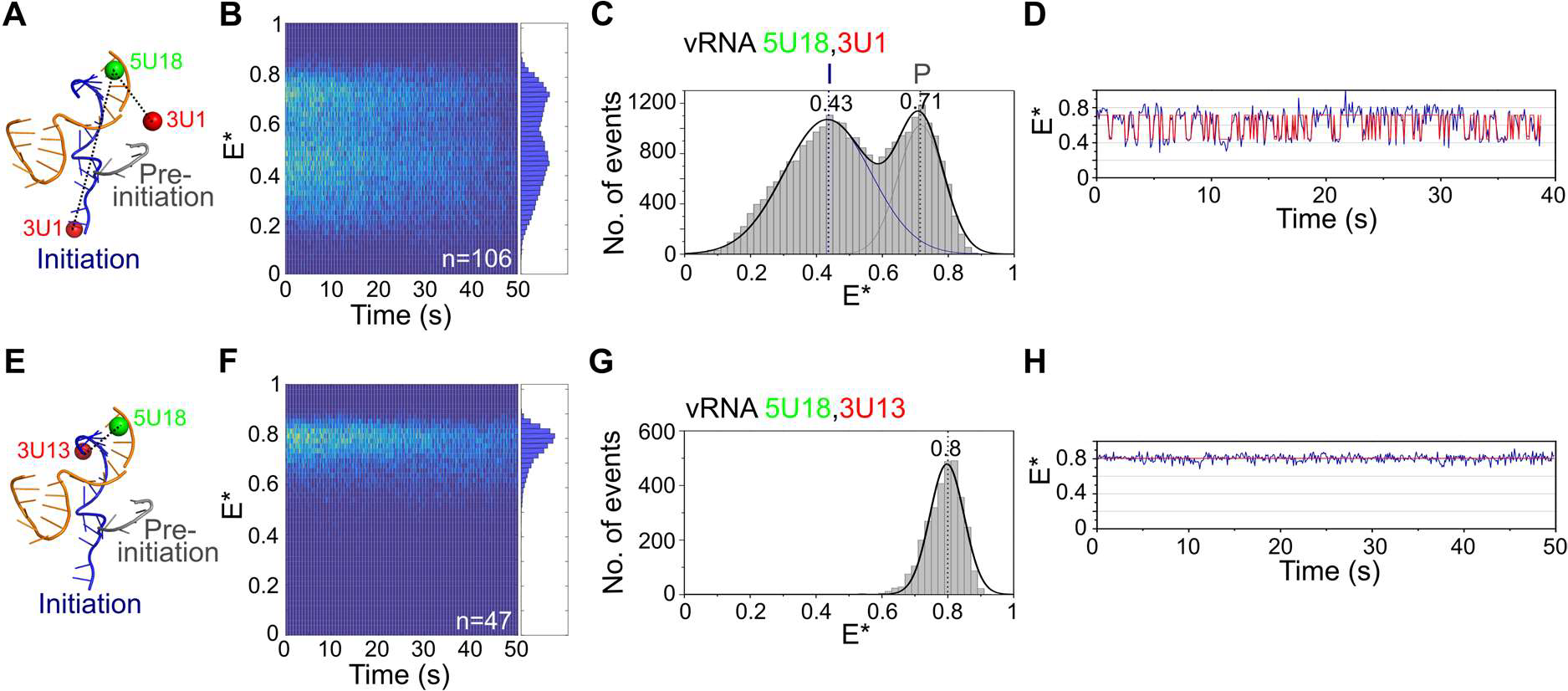
The 3’ vRNA in immobilized replication complexes is dynamic. A) Schematic of conformations of the vRNA promoter in the absence of nucleotides, labelled with donor and acceptor fluorophores at positions 18 on the 5’ end and 1 on the 3’ end. B) Kymograph of single-molecule time traces from immobilized replication complexes in the absence of nucleotides. E* represents apparent FRET efficiency. C) FRET histogram from B) fitted with a double Gaussian function. Initiation state = I, pre-initiation state = P. Frame time: 100ms. D) Example time trace showing transitions between the initiation and pre-initiation states. E) Schematic of the vRNA promoter labelled with donor and acceptor fluorophores at positions 18 on the 5’ end and 13 on the 3’ end. F) Kymograph of single-molecule time traces from immobilized replication complexes in the absence of nucleotides. G) FRET histogram from F). H) Example time trace showing a static FRET trace.

To verify that the FRET fluctuations were due to changes in the distance between the dyes (and hence due to the movement of the 3’ vRNA), we measured FRET between dyes located at positions 18 on the 5’ vRNA and 13 on the 3’ vRNA (Fig. 2E, S3B and S4B). Here, the dyes are located in the distal region of the vRNA promoter, a region that remains double-stranded and should not show FRET fluctuations. The dyes are close together in this configuration, and generated only a single high-FRET distribution centred at E*~0.80 (Fig. 2F&G). Individual time traces also showed a static FRET state, with no dynamics of the vRNA occurring in this region of the promoter (Fig. 2H). These results confirm that the dynamics in the absence of nucleotides were due to movement of the RNA within the proximal region of the vRNA promoter, while the distal region remained static.

We also examined the dynamics of replication complexes containing promoter vRNA labelled with donor and acceptor fluorophores at positions 18 on the 5’ strand and 4 on the 3’ strand (Fig. S6A), a labelling scheme that should also report on the conformational state of the 3’ vRNA. Similarly to the results obtained for the dye at position 1 on the 3’ strand, FRET histograms (Fig. S6B-D) and time traces (Fig. S6E) showed that the 3’ vRNA within individual complexes exhibited clear transitions between the initiation and pre-initiation states.

### Nucleotide addition stabilizes the 3’ vRNA initiation state

To study the effect of nucleotide addition on our immobilized replication complexes, we added ApG, a dinucleotide mimicking the initial product of de novo initiation, to the sample well prior to imaging (Fig. 3 and Fig. S7) (13). Similarly to the results obtained in the absence of nucleotides, we recovered a bimodal FRET distribution, centered at E*~0.34 for the initiation state and ~0.70 for the pre-initiation state (Fig. 3A). The shift of the low-FRET population to a lower E* value, compared to that measured when no ApG was present, is likely due to the increased distance of the dye at position 1 on the 3’ strand relative to the dye at position 18 on the 5’ strand, as the template RNA moves through the active site. ApG addition decreased the amplitude of the high-FRET population compared to when no ApG was present (34% to 12%), suggesting that nucleotide addition decreases the abundance of the pre-initiation conformation. We thus propose that the presence of the initiating nucleotide stabilizes the 3’ end of the vRNA in the active site. This observation was supported by individual time traces, which showed fewer transitions to the high FRET (pre-initiation) state (Fig. 3B).

**Figure 3.**
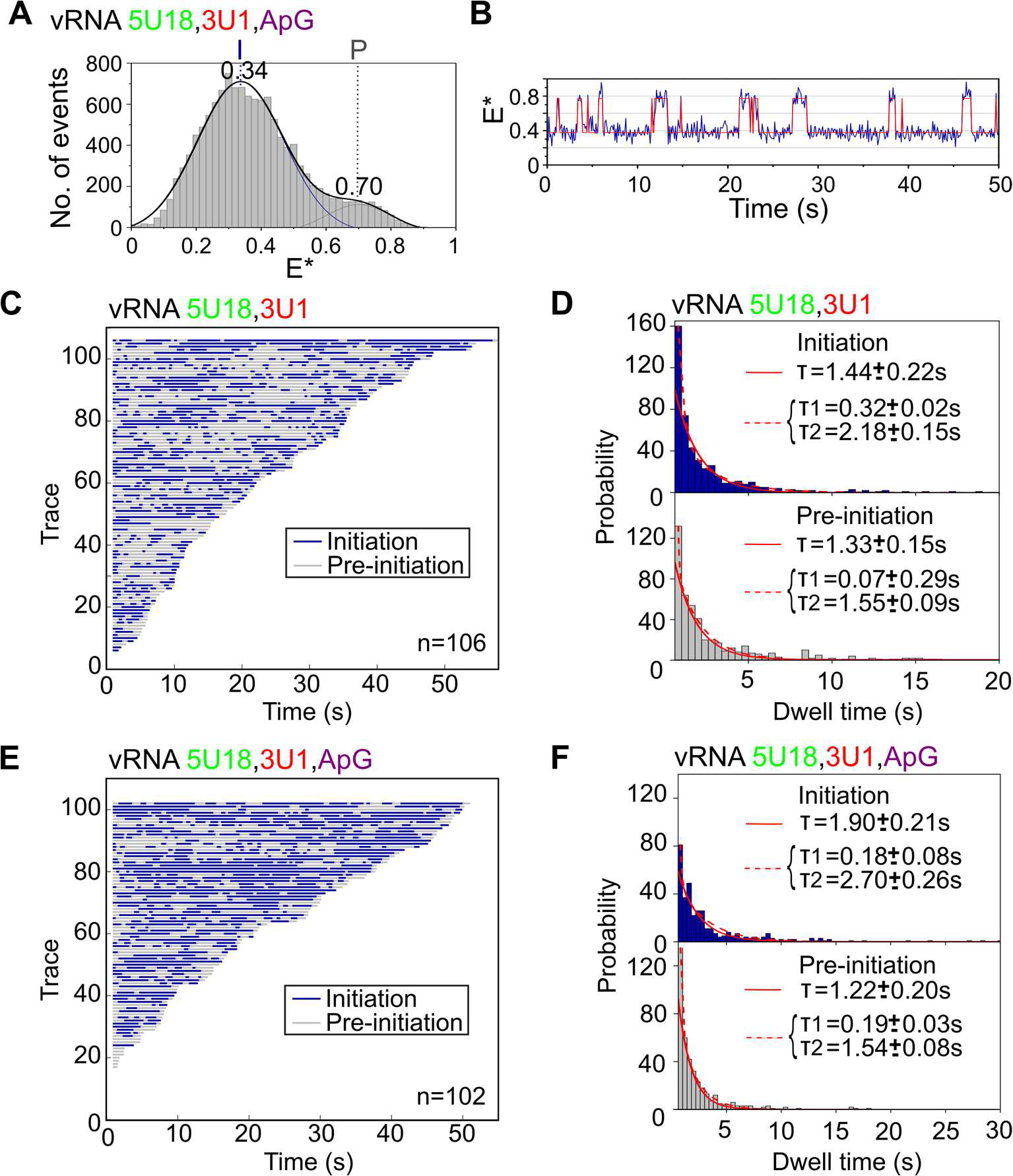
Nucleotide addition stabilizes the vRNA initiation state. A) Single-molecule FRET histogram from immobilized replication complexes after ApG addition. E* represents apparent FRET efficiency, initiation state = I, pre-initiation state = P, frame time: 100ms. B) Example time trace showing transitions between the initiation and pre-initiation states. C) Hidden Markov Modelling (HMM) was used to assign the states, which were colour-coded and binarised before being depicted in a single plot from shortest to longest in length. The low-FRET initiation state is represented in blue and the high-FRET pre-initiation state represented in grey. Number of traces (n) = 106. D) Histograms of initiation (top) and pre-initiation (bottom) dwell times for immobilized replication complexes in the absence of nucleotides. The data were fitted to a single-exponential function. E) Binarised time traces from immobilized replication complexes after 500µM ApG addition. Number of traces (n) = 102. F) Histograms of initiation (top) and pre-initiation (bottom) dwell times for immobilized replication complexes after ApG addition. The data were fitted to single (solid red line) or double-exponential (dotted red line) functions.

To further analyze the interconversion of the 3’ vRNA between the pre-initiation and initiation conformations, we used Hidden Markov Modelling (HMM) to assign the states. The two states in each trace were colour-coded, binarised, and sorted into a single plot from the shortest to longest in length. This analysis showed that, in the absence of nucleotides, some complexes showed rapid transitions between the initiation (shown in blue) and pre-initiation (shown in grey) states, while others showed slower transitions (Fig. 3C). To quantify the time that the vRNA spent in each conformational state, we obtained dwell-time distributions for the initiation and pre-initiation states. A single-exponential function did not fully take into account short dwells, and so the distributions were fitted to double-exponential functions, providing dwell times of 0.32±0.02 s and 2.18±0.15 s for the initiation state and 0.07±0.29 s and 1.55±0.09 s for the pre-initiation state (Fig. 3D). Upon ApG addition, individual complexes also showed a combination of rapid and slow transitions; however, replication complexes spent slightly more time in the initiation (blue) state (Fig. 3E), which was confirmed by a small increase in the longer dwell time for the initiation state (2.70±0.26 s) (Fig. 3F). We note that the large change in equilibrium seen in the FRET histogram when ApG was added is not reproduced by the modest increase in dwell times, perhaps due to a lack of equilibrium processes or multiple-exponential decay kinetics. However, these results are consistent overall with our previous solution-based smFRET observations showing that nucleotide addition stabilizes the 3’ vRNA initiation state and further, our results capture the timescale at which the 3’ vRNA accesses the initiation configuration.

### The cRNA promoter adopts multiple conformations but is not dynamic

To study the binding of the cRNA promoter by the influenza RNAP, we studied immobilized replication complexes containing short synthetic RNAs corresponding to the 5’ and 3’ termini of the cRNA. As structural data are only available for the first 12 residues of the 5′ cRNA (Fig. 4A) (12), we based our dye labelling positions on the vRNA promoter structure, using a donor dye placed at position 17 on the 5’ cRNA strand, and an acceptor dye at position 1 on the 3’ cRNA strand (Fig. 4B). We have shown previously that these positions did not affect the activity of the RNAP in an vitro replication assay (13).

**Figure 4.**
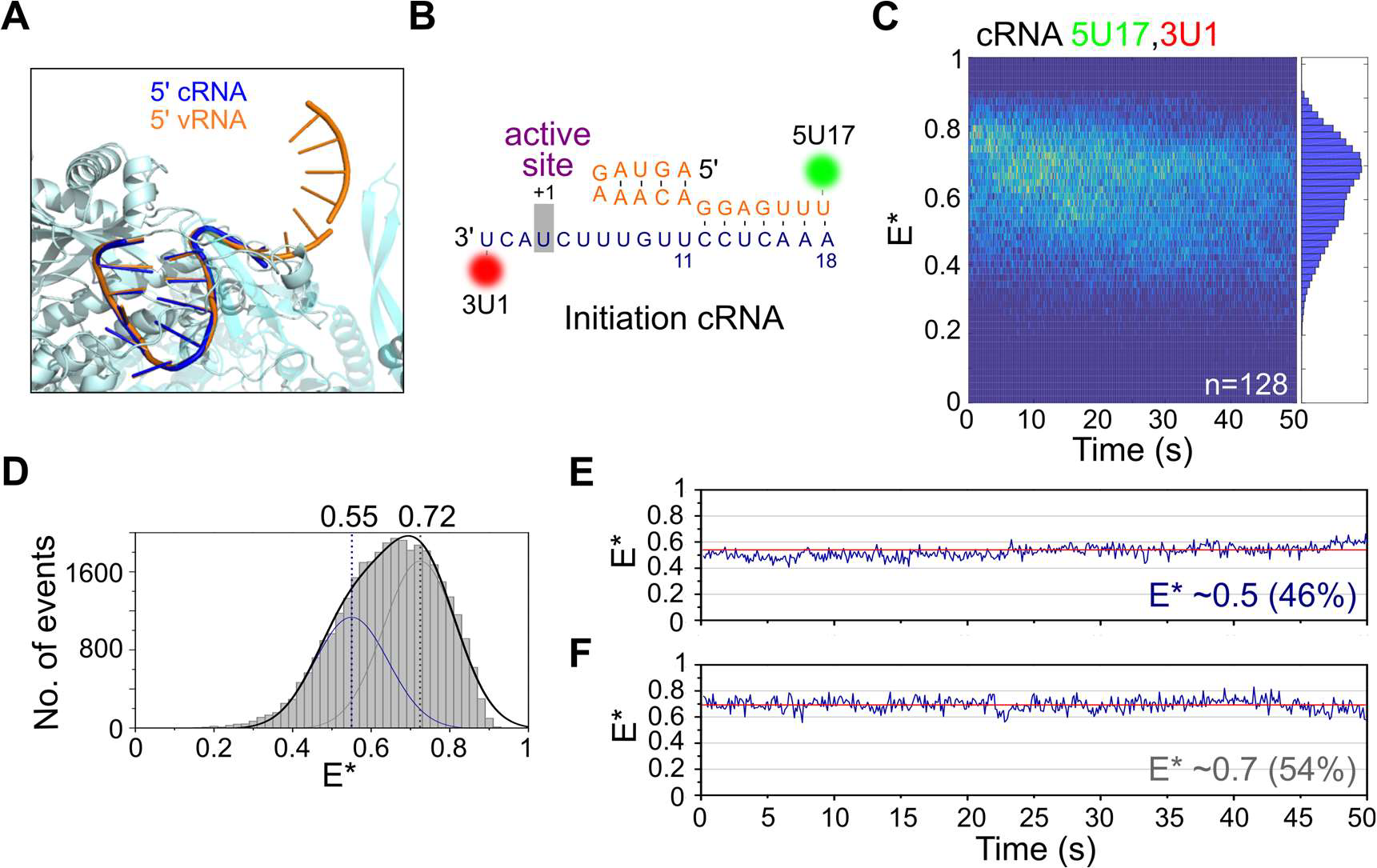
The cRNA promoter within immobilized replication complexes adopts multiple conformations but is not dynamic. A) The 5’ cRNA adopts a similar stem-loop structure to 5’ vRNA. 5’ vRNA (PDB code 5MSG) is shown in orange, 5’ cRNA (PDB code 5EPI) in dark blue, and RNAP (PDB code 5MSG) in cyan. B) Fluorophore positions and sequence of the cRNA promoter used in immobilized complexes. C) Kymograph of time traces from immobilized complexes containing a cRNA promoter labelled with a donor fluorophore at position 17 on the 5’ end and an acceptor fluorophore at position 1 on the 3’ end. E* represents apparent FRET efficiency. Frame time: 100ms. D) FRET histogram from C), fitted with Gaussian curves centred at 0.55 and 0.72. E&F) Representative time traces showing the lack of transitions between states.

We analyzed a large number (n=128) of replication complexes containing a labelled cRNA promoter and obtained a wide FRET distribution, corresponding to replication complexes with the cRNA promoter in multiple conformational states (Fig. 4C). The FRET distribution had a main peak at E*~0.7 and a shoulder at E*~0.5, and fitted well to a double Gaussian function with E*~0.55 and E*~0.72 (Fig. 4D). Based on the similarity between the vRNA and cRNA promoter sequences, we tentatively assigned the mid-FRET distribution to cRNA in the initiation state, and the high-FRET distribution to cRNA in the pre-initiation state.

Surprisingly, inspection of time traces from individual replication complexes revealed a striking difference between the vRNA and cRNA promoters. Unlike the vRNA promoter, in which the 3’ vRNA within individual complexes exhibited clear transitions between the initiation and pre-initiation states (Fig. 1D, Fig. 2D), the cRNA promoter did not show any conformational dynamics (Fig. 4E&F). Minor fluctuations in the static cRNA traces are likely due to stochastic photophysics of the dyes (26), and cRNA traces were never observed to show anti-correlated signals in the AA and DA intensities like vRNA traces (Fig S8). To exclude the possibility that transitions in the cRNA promoter occurred faster than our temporal resolution (~100 ms), we performed similar experiments at a frame time of 20 ms, but still did not observe any RNA dynamics (Fig. S9A). We also analyzed the FRET signal from immobilised cRNA replication complexes at 37°C, and found that, although the FRET traces became noisier, there were no transitions between states (Fig. S9B).

As a control for cRNA promoter dynamics, we used a similar duplex region labelling scheme as the one used for the vRNA promoter, and measured FRET on immobilised complexes containing cRNA labelled with donor and acceptor fluorophores at position 17 on the 5’ strand and position 14 on the 3’ strand, respectively. Similarly to the results from the vRNA promoter, we observed a single FRET distribution, centred at E*0.75 (Fig. S10A&B). Individual time traces also showed a static FRET state, with no dynamics of the cRNA occurring in this region of the promoter (Fig. S10C). Taken together, our FRET results suggest that, unlike the highly dynamic vRNA promoter, the cRNA promoter has multiple conformational states that do not interconvert significantly within the timescale of our experiments (~1 min).

### Nucleotide addition results in a stable low-FRET cRNA conformation

Unlike the vRNA promoter, for which replication initiates terminally at position 1 on the 3’ template (Fig. 5A), replication of the cRNA promoter is thought to initiate internally at positions 4 and 5 (hereafter called initiation complex ic(4,5)), followed by realignment of the dinucleotide product to bases 1 and 2 (hereafter called ic(1,2)) (Fig. 5B). To study the effect of nucleotide addition on the conformation of the cRNA promoter, we added ApG to immobilised replication complexes that contained promoter cRNA labelled at position 17 on the 5’ strand and 1 on the 3’ strand. Similarly to complexes in the absence of nucleotides, a wide FRET distribution was observed (Fig. 5C), indicative of multiple conformational states, and was fitted with Gaussians centred at E*~0.40, E*~0.55 and E*~0.72 (Fig. 5D). Based on our results from the cRNA promoter conformation in the absence of nucleotides, we assigned the E*~0.55 population to cRNA initiation complexes, and the E*~0.72 population to pre-initiation complexes. Addition of ApG therefore results in the formation of a new, low-FRET state (E*~0.40), which likely represents ic(4,5) replication complexes wherein the 3’ cRNA template has translocated further into the active site to allow internal initiation at position 4.

**Figure 5.**
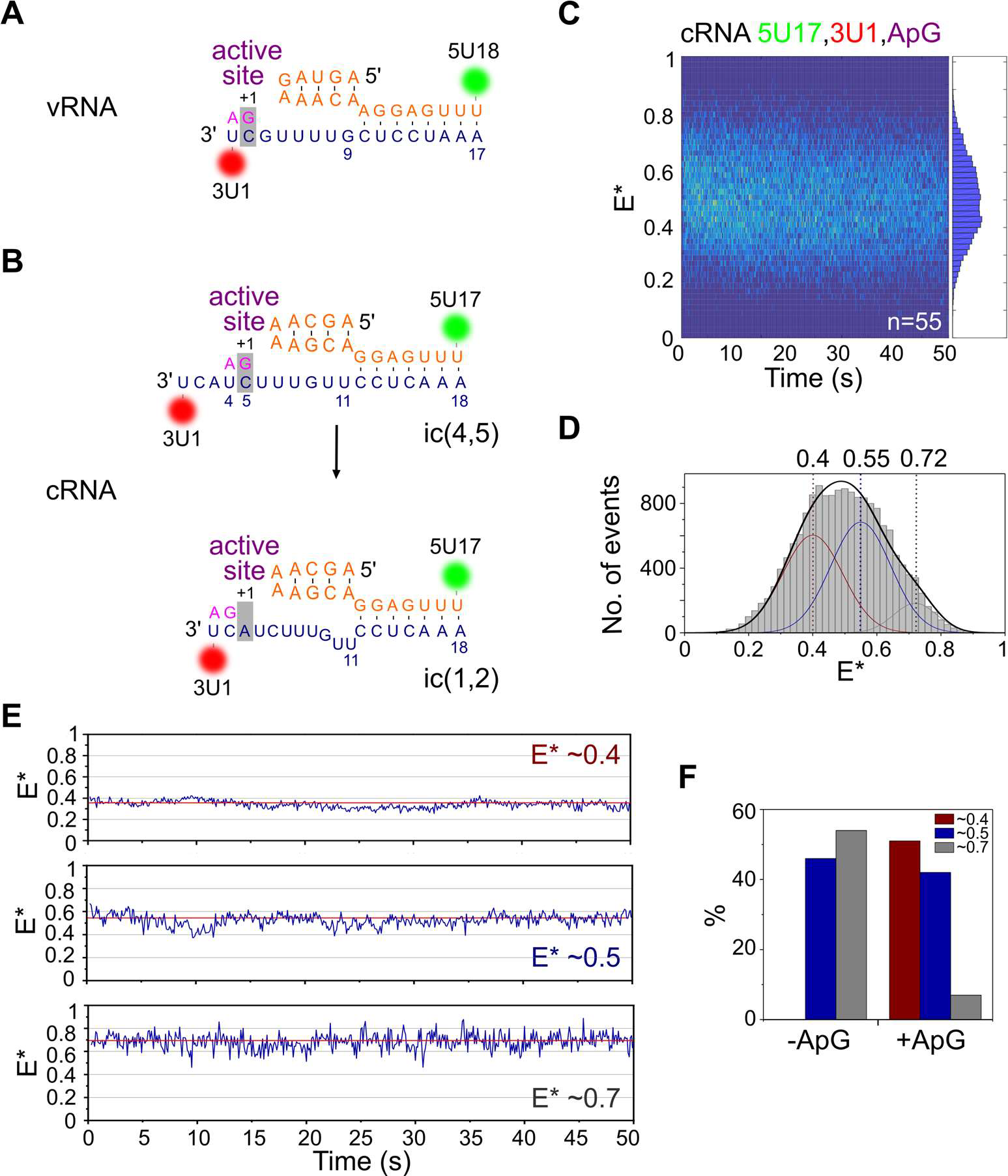
Nucleotide addition results in a stable low-FRET state for the cRNA promoter. A) Schematic of replication initiation on the vRNA promoter, which initiates terminally at position 1 on the 3’ template. B) Schematic of the prime-and-realign mechanism of replication initiation on the cRNA promoter. Replication initiates internally, from position 4 on the 3’ template (initiation complex(4,5)), followed by realignment of the nascent dinucleotide vRNA product to position 1 (ic(1,2)). C) Kymograph of time traces from immobilized complexes containing a cRNA promoter labelled with donor and acceptor fluorophores at position 17 on the 5’ end and position 1 on the 3’ end respectively, after 500µM ApG addition. E* represents apparent FRET efficiency. Frame time: 100ms. D) FRET histogram from C), fitted with Gaussian curves centred at 0.40, 0.55 and 0.72. Peaks centred at mean E* 0.55 and 0.72 were fixed, while the peak centred at E* 0.40 was fitted freely. E) Representative time traces showing the lack of transitions between states. F) Time traces were classified into groups and compared in the absence (-ApG) or presence (+ApG) of nucleotides.

Analysis of time traces from individual replication complexes revealed that the cRNA promoter did not show any conformational dynamics upon ApG addition, with the vast majority of molecules (99%) showing no transitions between states (Fig. 5E). Time traces were manually classified into three groups representing complexes in which the cRNA was in a static low-FRET conformation (E*~0.4), static mid-FRET conformation (E*~0.5), or static high-FRET conformation (E*~0.7). We calculated the fraction of replication complexes in each of these conformations, and compared it to the conformations obtained without nucleotides. When ApG was absent, no low-FRET complexes were observed, cRNA in 46% of the complexes was in a mid-FRET state (E*~0.5; corresponding to the initiation state) and cRNA in 54% of the complexes was in a high-FRET state (E*~0.7; corresponding to the pre-initiation state) (Fig. 5F and Fig. 4E&F). By contrast, when ApG was present, half (~51%) of the complexes formed a stable low-FRET state (E*~0.4; consistent with ic(4,5) translocated complexes), with the remaining complexes being either in a mid-FRET state (42%; initiation state) or high-FRET state (7%, pre-initiation state) (Fig. 5E). Unlike the vRNA promoter, where the 3’ vRNA remained dynamic upon nucleotide addition but spent more time in the active site of the RNAP, the cRNA pre-initiation state largely disappeared upon nucleotide addition, suggesting that a stable translocated state is formed upon nucleotide addition. Our results highlight large differences in the initiation mechanisms for the vRNA and cRNA promoter, in agreement with previous reports (14, 15) suggesting that replication initiates terminally for vRNA, and internally for cRNA.

### Additional residues in the proximal 3’ cRNA are responsible for stable promoter binding

To dissect the structural determinants of the observed differences in vRNA and cRNA promoter binding dynamics, we first examined the role of the priming loop, a β-hairpin in the PB1 subunit that acts as a stacking platform for the initiating base during terminal initiation on the 3′ vRNA (15). Since we hypothesize that the priming loop may stabilize the cRNA 3’ strand in the active site, we deleted residues 642-656 of the PB1 subunit of the RNAP, corresponding to the full priming loop (Fig. S11A) (16); the resulting mutant RNAP was previously shown to have a higher activity than wild type RNAP in an in vitro capped primer extension assay (16).

To examine the effect of the priming loop on the conformations of promoter RNA, we measured FRET on immobilised complexes containing Δ642-656 RNAP and either vRNA labelled at position 18 on the 5’ end and 1 on the 3’ end, or cRNA labelled at position 17 on the 5’ end and 1 on the 3’ end. For Δ642-656 RNAP-vRNA complexes, the two FRET populations were not as distinct as those observed with wildtype RNAP, giving rise to a narrower overall FRET distribution (Fig. S11B, top); however, inspection of individual time traces showed that, as with wildtype RNAP, the vRNA promoter was highly dynamic, with clear transitions between initiation and pre-initiation states (Fig. S11C, top). The overall FRET distribution for the cRNA promoter was bimodal like wildtype, however, unlike wildtype, some time traces showed transitions between states (Fig. S11B and S11C, bottom). Comparison of the wildtype and Δ642-656 RNAP complexes exhibiting dynamics showed that priming loop deletion did not change the number of dynamic vRNA complexes (87% for both wildtype and Δ642-656 RNAP) (Fig. S11D). However, there was an increase in the number of Δ642-656 RNAP-cRNA complexes that showed dynamics (11% compared to 1% using wildtype RNAP) (Fig. S11D). These results suggest that the priming loop has some effect on cRNA 3’ end dynamics, but the loop deletion is not sufficient to bring the number of dynamic traces near to vRNA promoter levels.

Since the promoter RNA sequence is the only difference between vRNA and cRNA promoter complexes, we turned our attention to the sequence of the 3’ cRNA strand. The sequence of the proximal 3’ cRNA strand before the distal double-stranded duplex region is two nucleotides longer than that of the proximal 3’ vRNA strand (cf. Fig. 5A&B), possibly because during cRNA initiation the 3΄ template must overreach the active site cavity to allow internal initiation at position 4. We hypothesized that the longer length of the cRNA 3’ strand, compared to the vRNA 3’ strand, was responsible for the lack of cRNA dynamics observed in our earlier experiments. We therefore deleted residues 6 and 7 of the 3’ cRNA (Fig. 6A), and found that truncation of the 3’ strand did not affect the ability of the cRNA promoter to bind RNAP (Fig. 6B). Similarly to the results obtained using a wildype cRNA promoter, the FRET histogram from immobilized complexes containing RNAP and Δ6,7 cRNA was bimodal (Fig. 6C). This was fitted with a double Gaussian function, centred at E*~0.43 and E*~0.66. When we analyzed time traces from individual complexes we found that, while some traces (39%) remained static (Fig. 6D), most complexes (61%) showed transitions between the initiation and pre-initiation states (Fig. 6E). This supports our suggestion that the additional residues of the proximal 3’ cRNA, compared to the 3’ vRNA, limit dynamics and promote the stable binding of the cRNA by the RNAP. Intriguingly, these additional two residues are also crucial for initiation in a cellular context (Fig. S12), likely because they are necessary for internal initiation to occur.

**Figure 6.**
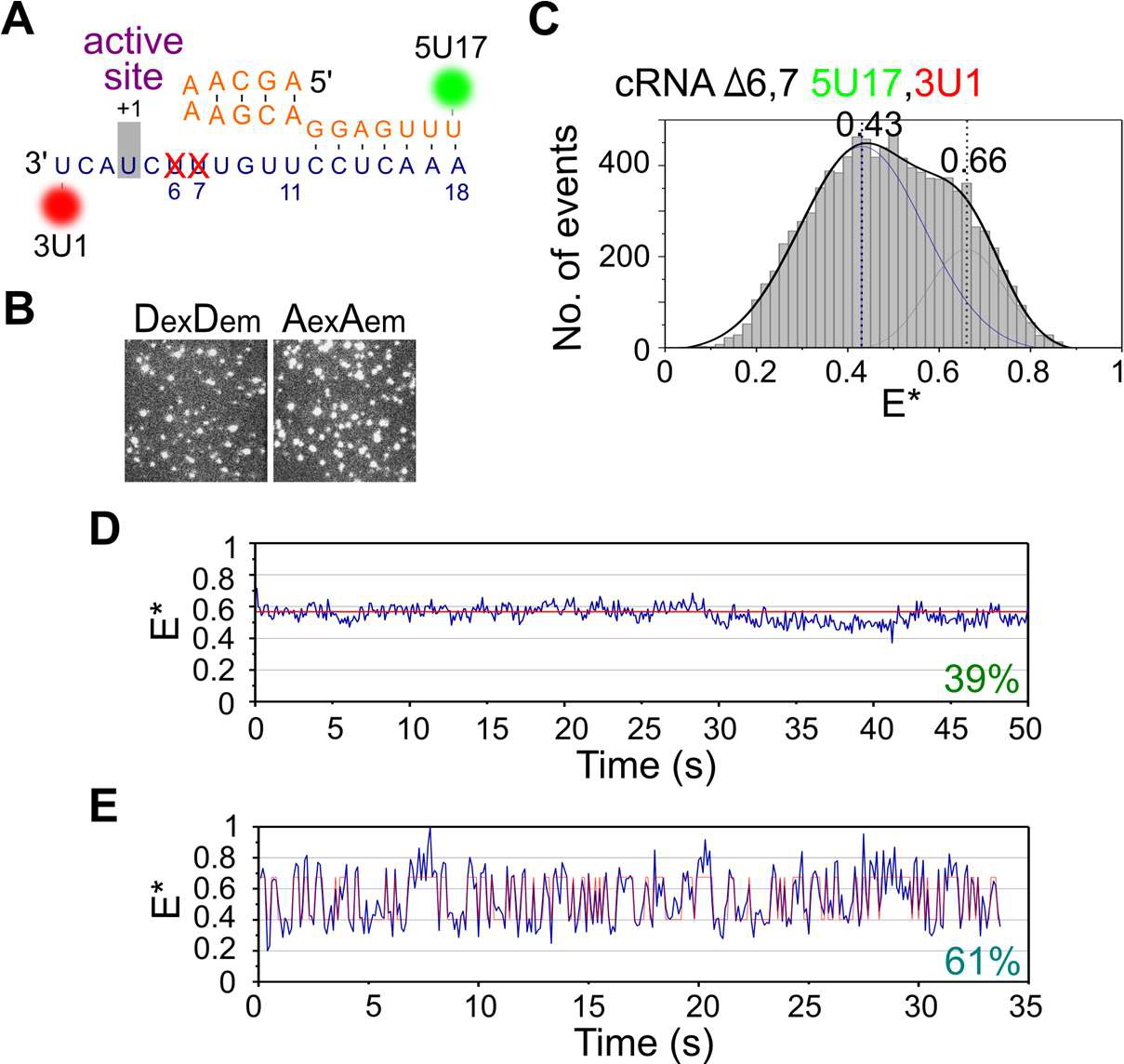
Truncation of the 3’ cRNA restores dynamics. A) Schematic of the cRNA promoter, showing deletion of residues 6 and 7 of the 3’ strand. B) Representative field-of-view of the donor (DexDem) and acceptor (AexAem) channels from immobilized replication complexes containing a Δ6,7 cRNA promoter labelled with donor and acceptor fluorophores at position 17 on the 5’ end and position 1 on the 3’ end respectively, showing that deletion of residues 6 and 7 does not hinder promoter binding. C) FRET histogram from B), fitted with Gaussian curves centred at 0.43 and 0.66. E* represents apparent FRET efficiency. Frame time: 100ms. D&E) Representative time traces of immobilized complexes.

## DISCUSSION

Structural and biophysical studies have demonstrated that the RNA of the influenza virus can adopt multiple conformational states; however, the dynamics of these states and the factors that influence them are unknown. Here, we observe the dynamic motions of promoter RNA in real time using single immobilized replication complexes. We discovered distinct differences in dynamics between short synthetic 3’ vRNA and cRNA templates, highlighting differences in the initiation mechanisms between the two promoters. We expect that the observed dynamics will also be present in replication complexes with longer genomic RNA segments, although the time constants of dynamics may be affected by longer RNA. The dynamics of the 3’ vRNA are thermally driven, since transitions were observed in the absence of any nucleotides, i.e., no ATP consumption was required for positioning the 3’ vRNA in the RNAP active site.

The pre-initiation state, thus far only observed in structures of the RNAP containing the vRNA promoter (4), has been suggested to represent an inactive state of the RNP, possibly the vRNA conformation found within vRNP complexes packaged into progeny virions, which are not actively replicating. This hypothesis is supported by our finding that ApG addition stabilises the RNAP complex in an initiation state, which presumably represents the active form of the RNA within RNAP. Our results also show that in most complexes, the vRNA promoter is highly dynamic, exhibiting rapid transitions between different states. vRNA dynamics may therefore be a requirement for robust RNAP activity, as vRNA initiation complexes have been found to be ~200-fold more efficient in transcription initiation than cRNA initiation complexes (6). Intriguingly, this suggests that the rapid dynamics of the 3’ vRNA may be the key to fast initiation, possibly because the dynamic nature of the vRNA promoter gives the 3’ RNA a greater chance of accessing the active site, and thus of being stabilized if nucleotides are present.

Our previous work using solution-based smFRET on replication complexes revealed that the addition of RNAP to the cRNA promoter resulted in a wide FRET distribution, indicative of multiple RNA conformations (13). We interpreted this to mean that the cRNA promoter adopted a similar structure to the vRNA promoter, with initiation and pre-initiation states. Analysis of immobilised cRNA complexes confirmed this, although examination of individual molecules revealed that the 3’ cRNA was stably bound in each conformation, rather than being dynamic. The conferred stability could be due to RNA-RNA interactions, RNAP–RNA interactions or RNAP conformational changes. Indeed, crystal structures suggest that there are major conformational differences in the arrangement of peripheral domains of the RNAP between the apo, vRNA-bound, or cRNA-bound forms of the RNAP (12), and the priming loop has been shown to be important for stabilizing terminal initiation, but not internal initiation (15). Deletion of the full priming loop in our experiments resulted in a small number of cRNA initiation complexes showing 3’ dynamics (11%), suggesting that the priming loop may play a minor role in the stabilization of the cRNA 3’ binding.

Crystal structures of the RNAP have shown that the vRNA 3’ end overshoots the active site by 1 nucleotide, meaning that for terminal initiation on the vRNA template to occur, the RNA needs to move back by one nucleotide (Fig. 5A) (6). Sequence comparison of the vRNA and cRNA promoters shows that the proximal 3’ cRNA is one nucleotide longer than the 3’ vRNA, and the distal promoter duplex region is one nucleotide shorter (compare Fig. 5A and B). This suggests that the cRNA promoter has an alternative duplex region base-pairing scheme to that of vRNA promoter, involving 3’ nucleotides 12-14 (instead of 10–12 as in the vRNA) with 5΄ nt 11–13. This would make the cRNA 3’ strand overshoot the active site by three nucleotides, ideally positioning it for internal initiation from nucleotide 4 (Fig. 5B). We showed that truncation of the 3’ cRNA by two bases was sufficient for 61% of the complexes to exhibit dynamics similar to those observed for the vRNA promoter. The longer length of the proximal 3’ cRNA strand, compared to the proximal 3’ vRNA strand, therefore stabilizes the RNA in the RNAP active site. Stable binding of the cRNA 3’ end in the active site, compared to the dynamic motions of the vRNA 3’ end, may be important for efficient internal initiation and the ‘prime-and-realign’ mechanism, which is crucial for synthesis of a full-length genomic vRNA (16). Similar strategies may exist in members of other virus families, such as the arenaviruses and bunyaviruses (34, 35), which also use prime-and-realign mechanisms for the initiation of RNA synthesis.

In summary, our results advance our understanding of influenza virus replication by providing novel insights into the differences in initiation mechanisms for the vRNA and cRNA promoters. Real-time monitoring of promoter binding at the level of single RNAP complexes has revealed mechanistic details unavailable from ensemble transcription assays or static crystal structures, and has opened an avenue to study transitional intermediates as they occur along the initiation pathway. Our methods should also be able to examine other, alternative promoter RNA conformations (such as the extended, single-stranded configuration of the proximal 3’ RNA observed in the flu-related La Crosse bunyavirus RNAP structure (10)) and our findings should be widely applicable to the study of replication in other viruses.

