## Supplemental files for "Real-time analysis of single influenza virus replication complexes reveals large promoter-dependent differences in initiation dynamics"

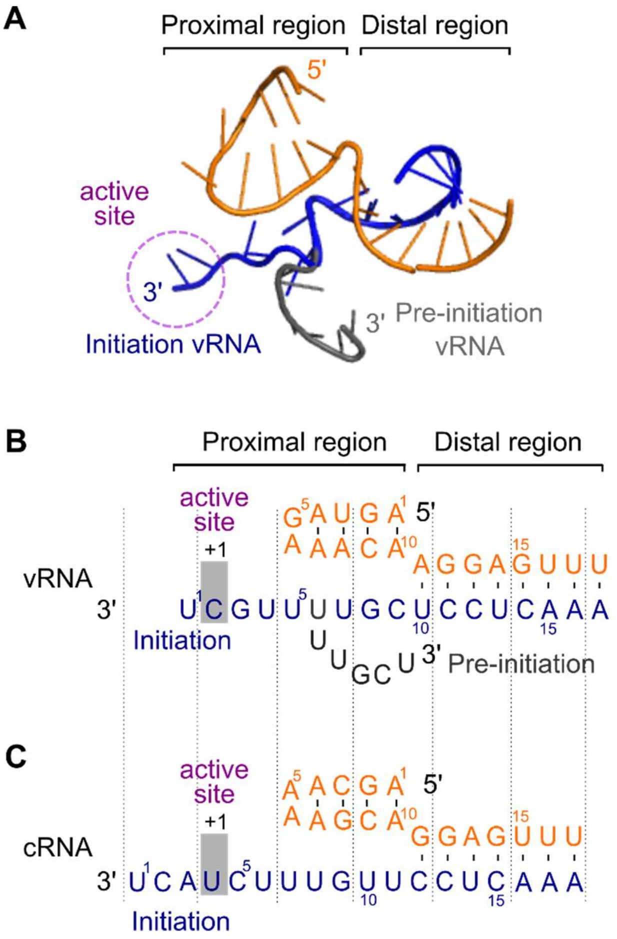


**Supplementary Figure 1. Comparison of vRNA and cRNA promoter sequences.** A) Conformations of 5ˈ vRNA and 3ˈ pre-initiation and initiation vRNA from influenza RNAP structures (1, 2). B) Sequence and predicted conformation of the vRNA promoter. C) Sequence and predicted conformation of the cRNA promoter.

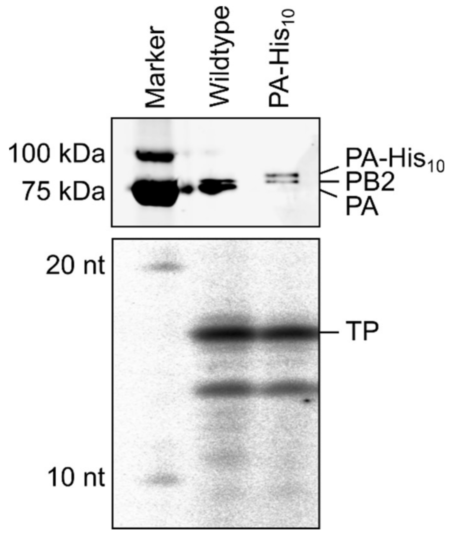


**Supplementary Figure 2. Purified histidine-tagged RNAP is active in vitro.** Wildtype or histidine-tagged RNAP (with a deca-histidine tag and a protein-A tag on the C-terminus of the PB2 subunit) were expressed in mammalian cells and purified using IgG-sepharose. A western blot using anti-PA and anti-PB2 antibodies confirmed the presence of the histidine tag (top panel). An in vitro assay showed no difference in transcriptional activity between wildtype and his-tagged RNAP using a synthetic RNA promoter as a template (lower panel). The full-length 17 nt-long transcription product is indicated (TP); shorter products represent premature termination or degradation products.

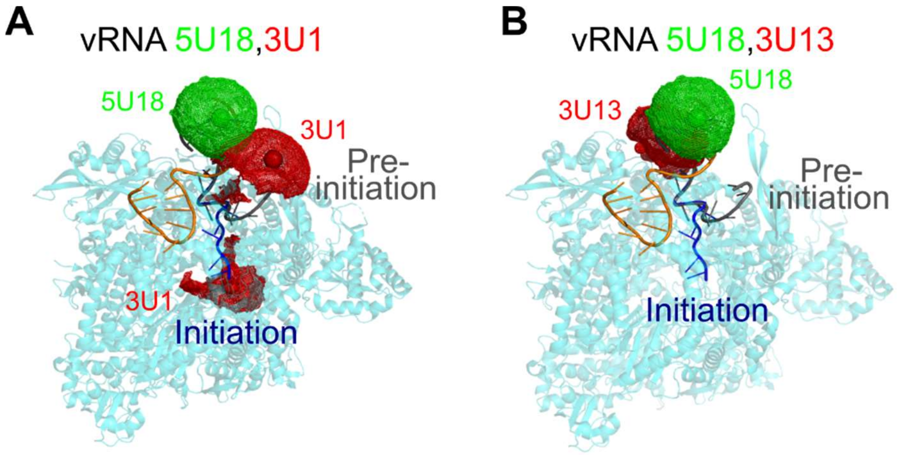


**Supplementary Figure 3. vRNA fluorophore positions and accessible volume modelling.** A) Average fluorophore positions (red or green spheres) and dye clouds corresponding to dye accessible volumes for the vRNA promoter labelled with donor and acceptor fluorophores at positions 18 on the 5' end and 1 on the 3' end. B) Same as A) except for fluorophore positions 18 on the 5' end and 13 on the 3' end.

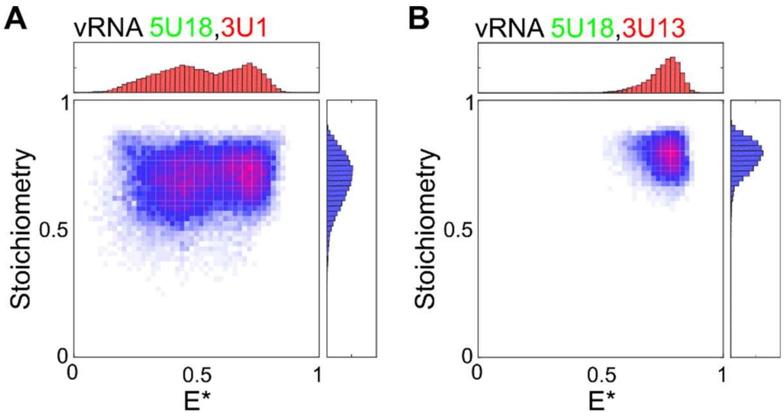


**Supplementary Figure 4. The 3ˈ vRNA in immobilized replication complexes is dynamic.** A) Single-molecule FRET histogram from immobilized replication complexes containing promoter vRNA labelled with donor and acceptor fluorophores at positions 18 on the 5' end and 1 on the 3' end. E* represents apparent FRET efficiency, frame time: 100ms. B) As in A), but with promoter vRNA labelled with donor and acceptor fluorophores at positions 18 on the 5' end and 13 on the 3' end.

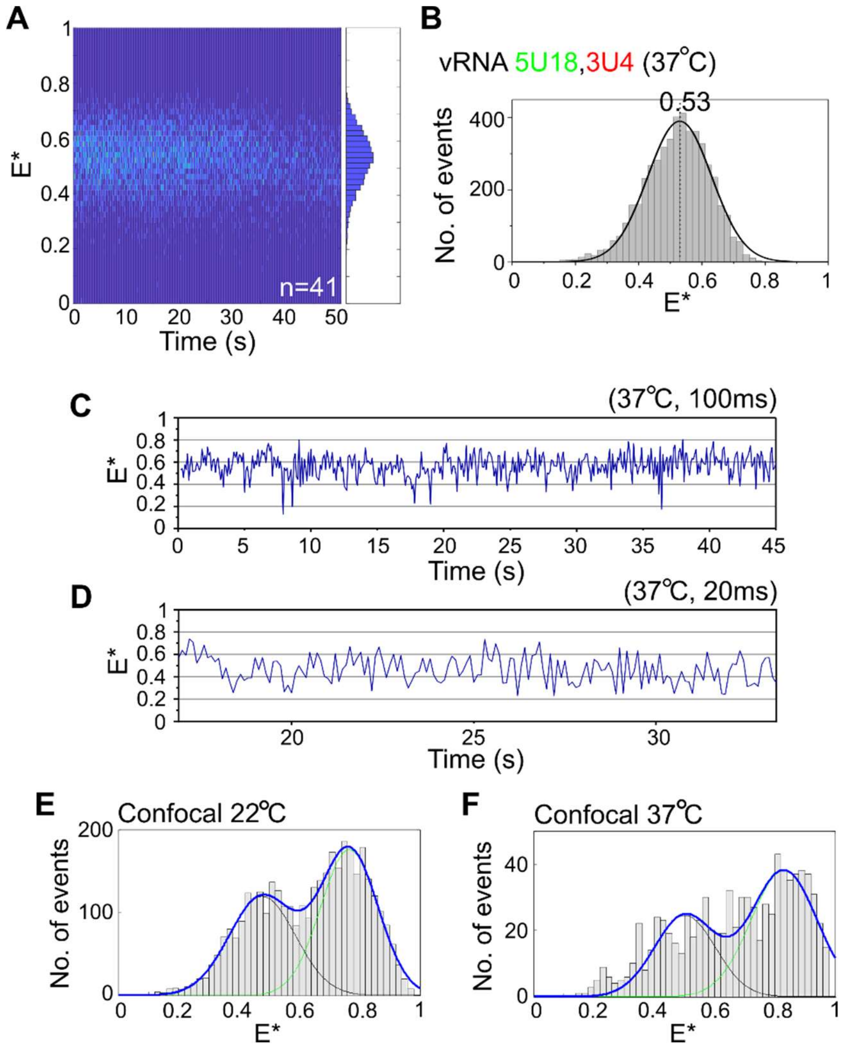


**Supplementary Figure 5. Analysis of the vRNA promoter within replication complexes at higher temperature.** A) Kymograph of time traces from immobilized complexes containing a vRNA promoter labelled with donor and acceptor fluorophores at position 18 on the 5ˈ end and position 1 on the 3ˈ end respectively, acquired at 37°C. E* represents apparent FRET efficiency. Frame time: 100ms. B) FRET histogram from A), fitted with a Gaussian curve centred at 0.53. C) Representative time trace at a frame time of 100ms. D) Representative time trace at a frame time of 20ms. E) FRET histogram from diffusing complexes, taken at 22°C. F) FRET histogram from diffusing complexes, taken at 37°C.

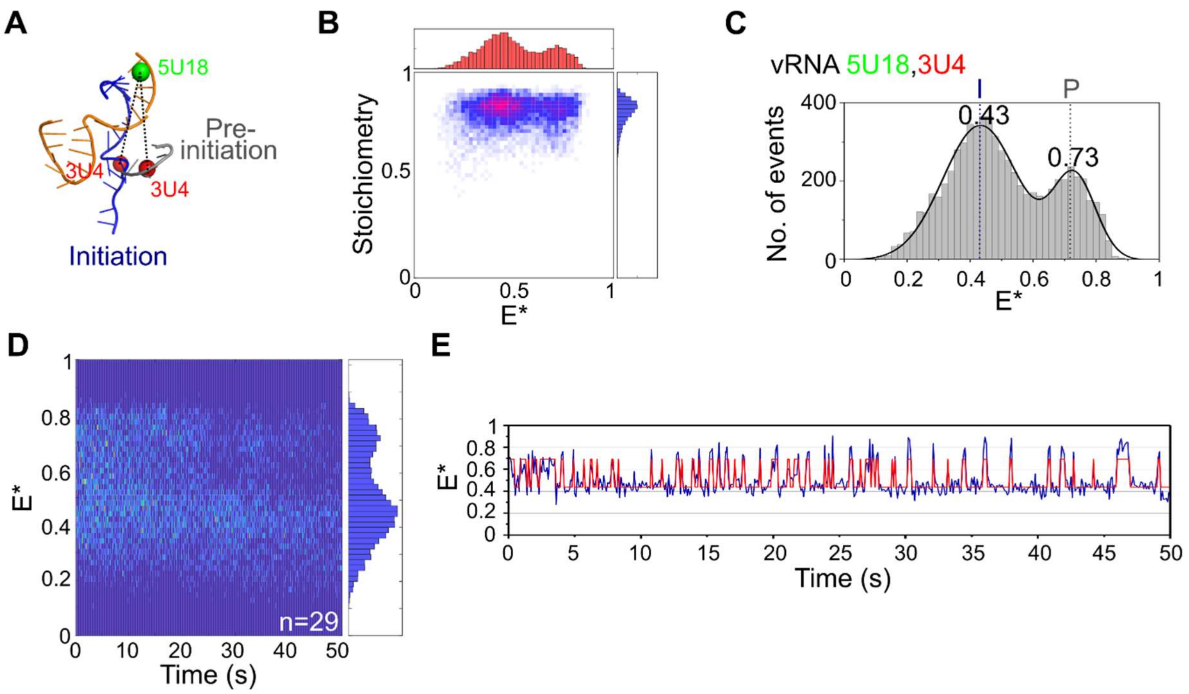


**Supplementary Figure 6. The vRNA promoter labelled at position 4 on the 3ˈ RNA adopts multiple conformations.** A) Schematic of conformations of the vRNA promoter, labelled with donor and acceptor fluorophores at positions 18 on the 5' end and 4 on the 3' end. B) ES histogram from immobilized replication complexes, where E* represents apparent FRET efficiency. Frame time: 100ms. C) FRET histogram from immobilized replication complexes, initiation state = I, pre-initiation state = P. D) Kymograph of time traces from immobilized vRNA complexes. E) Example time trace showing transitions between the initiation and pre-initiation states.

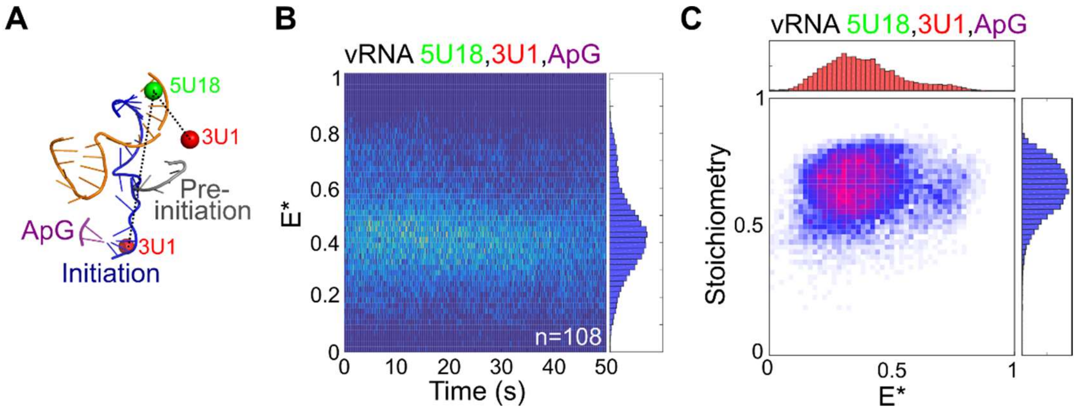


**Supplementary Figure 7. Nucleotide addition stabilizes the vRNA initiation state.** A) Schematic of conformations of the vRNA promoter in the presence of 500µM of the dinucleotide ApG. B) Kymograph of time traces from immobilized vRNA complexes after ApG addition, where E* represents apparent FRET efficiency. Frame time: 100ms. C) Single-molecule FRET histogram from immobilized replication complexes after ApG addition.

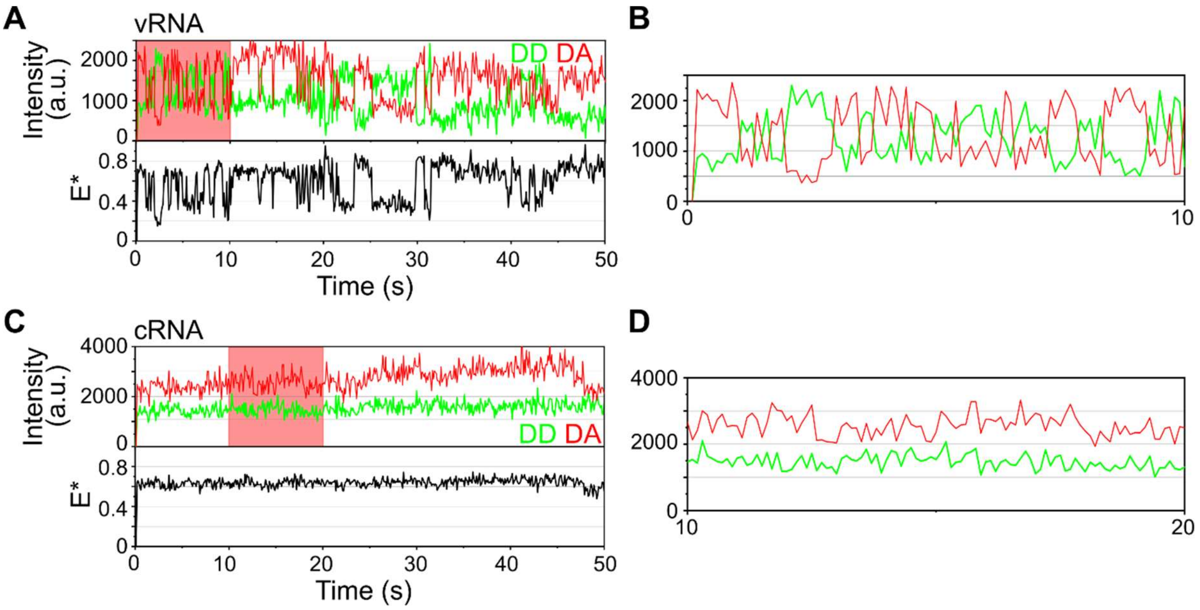


**Supplementary Figure 8. Comparison of donor and acceptor intensity signals between vRNA and cRNA.** A) Representative time trace from immobilized complexes containing a vRNA promoter labelled with donor and acceptor fluorophores at position 18 on the 5ˈ end and position 1 on the 3ˈ end respectively. E* represents apparent FRET efficiency. Frame time: 100ms. B) Zoom in on donor (DD)-acceptor (AA) anti-correlations from the region highlighted in red in A). C&D) Same as A&B except for the cRNA promoter labelled with donor and acceptor fluorophores at position 17 on the 5ˈ end and position 1 on the 3ˈ end respectively.

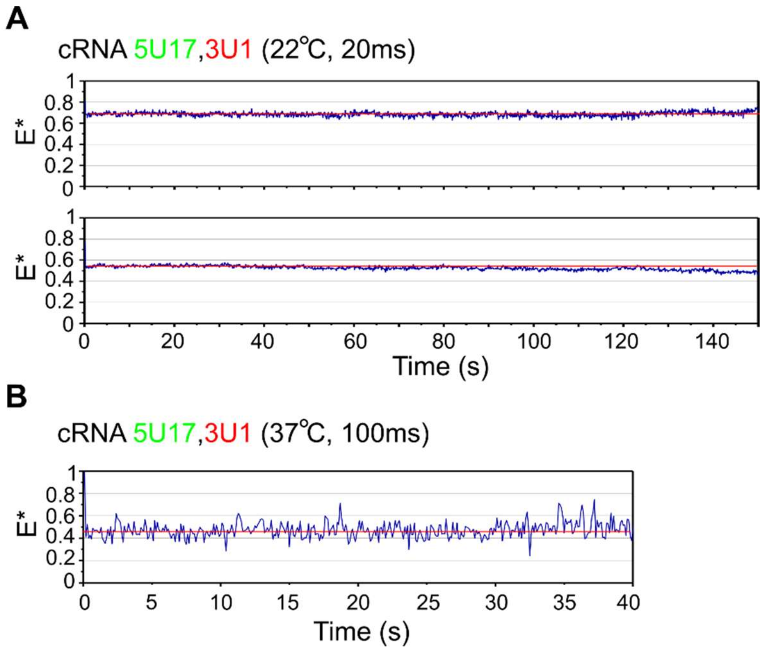


**Supplementary Figure 9. The cRNA promoter within immobilized replication complexes is not dynamic at a greater temporal resolution and higher temperature.** A) Representative time traces from immobilized complexes containing cRNA labelled with donor and acceptor fluorophores at position 17 on the 5ˈ end and position 1 on the 3ˈ end respectively. E* represents apparent FRET efficiency. Temperature: 22°C. Frame time: 20ms. B) As in A), but with a temperature of 37°C and frame time of 100ms.

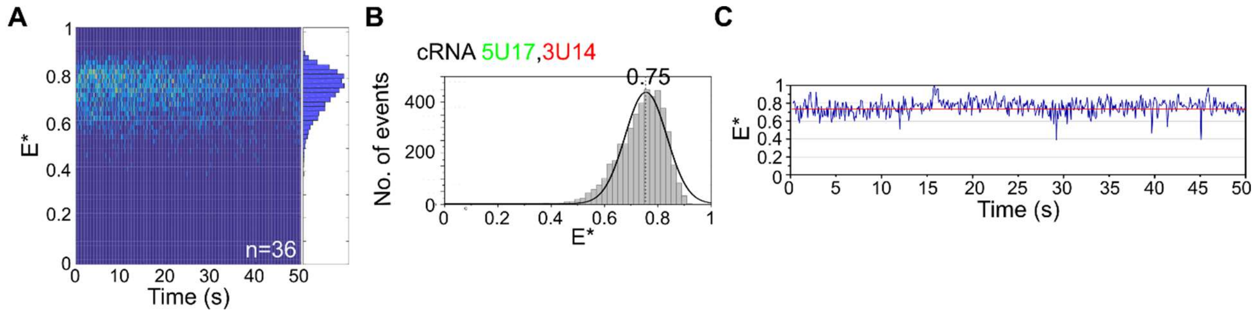


**Supplementary Figure 10. The duplex region of the cRNA promoter within immobilized replication complexes is not dynamic.** A) Kymograph of time traces from immobilized complexes containing RNA labelled with donor and acceptor fluorophores at position 17 on the 5ˈ end and position 14 on the 3ˈ end respectively. E* represents apparent FRET efficiency. Frame time: 100ms. B) FRET histogram from A). C) Representative time trace showing a static FRET signal.

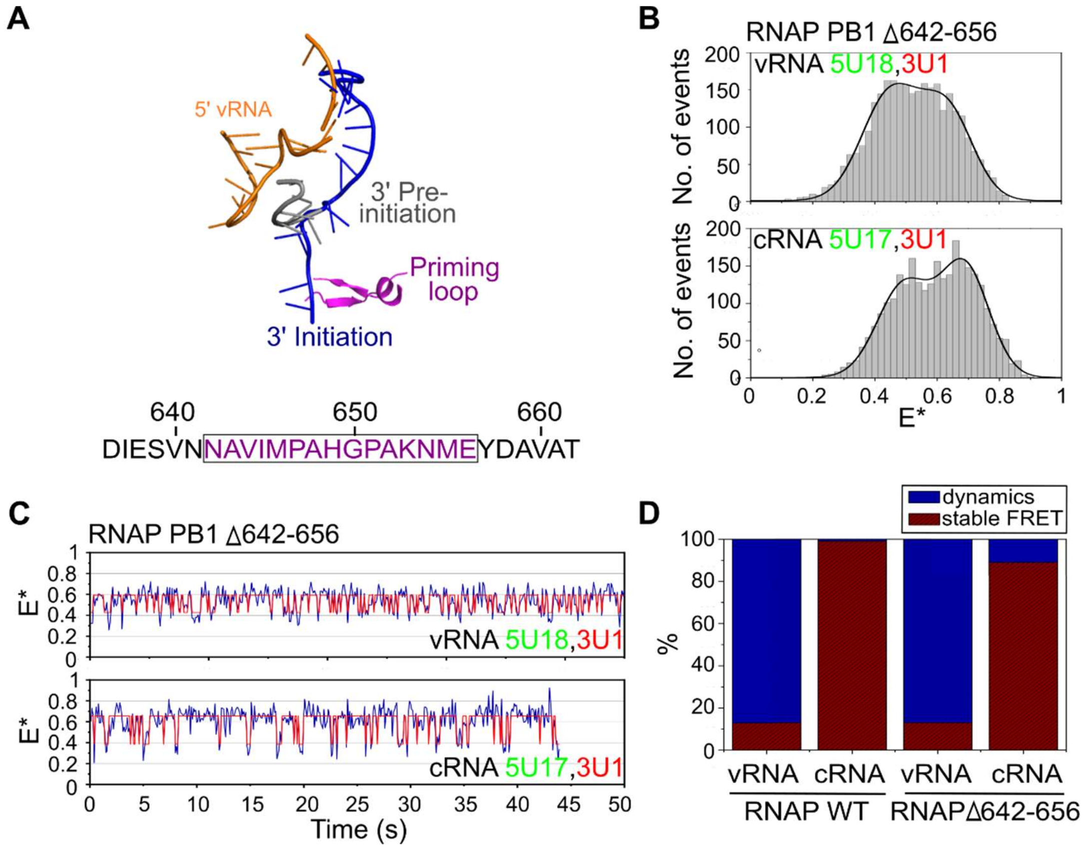


**Supplementary Figure 11. Deletion of the RNAP priming loop restores some cRNA dynamics.** A) Schematic showing the priming loop protruding towards the active site of the RNAP, where it stacks against the initiating NTP during vRNA to cRNA synthesis. Residues 642-656 of the PB1 subunit of the RNAP, corresponding to the full priming loop, were deleted. B) FRET histograms from immobilized complexes containing RNAP Δ642-656 and either vRNA labelled with donor and acceptor fluorophores at position 18 on the 5ˈ end and position 1 on the 3ˈ end, or cRNA labelled with donor and acceptor fluorophores at position 17 on the 5ˈ end and position 1 on the 3ˈ end. E* represents apparent FRET efficiency. Frame time: 100ms. C) Representative time traces showing transitions between states. D) Comparison of the % of molecules showing dynamics between the vRNA and cRNA promoters, using either wild-type (WT) or Δ642-656 RNAP.

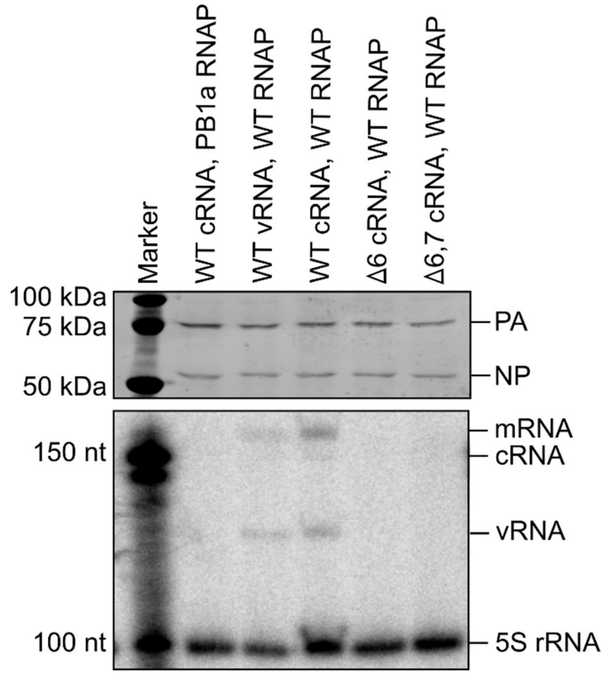


**Supplementary Figure 12. The Δ6,7 cRNA promoter has no activity in an RNP reconstitution assay.** Wildtype (WT) or active site mutant (PB1a) RNAP and either WT vRNA, or WT, Δ6 or Δ6,7 cRNA (segment 6) were expressed in mammalian cells. A western blot using anti-PA and anti-NP antibodies confirmed RNAP expression (top panel) and viral RNA levels (mRNA, cRNA and vRNA) were assessed by primer extension (lower panel). 5S ribosomal RNA (5S rRNA) levels were assessed as a loading control.
